# Genetic, textual, and archaeological evidence of the historical global spread of cowpea (*Vigna unguiculata* [L.] Walp)

**DOI:** 10.1101/2020.07.08.193995

**Authors:** Ira A. Herniter, María Muñoz-Amatriaín, Timothy J. Close

**Affiliations:** Department of Botany and Plant Sciences, University of California, Riverside, CA; Department of Plant Biology, Rutgers University, NJ; Department of Soil and Crop Sciences, Colorado State University, Fort Collins, CO

**Keywords:** *Vigna unguiculata*, domestication, archaeobotany, textual analysis

## Abstract

Cowpea (*Vigna unguiculata* [L.] Walp.) was originally domesticated in sub-Saharan Africa but is now cultivated on every continent except Antarctica. Utilizing archaeological, textual, and genetic resources, the spread of cultivated cowpea has been reconstructed. Cowpea was domesticated in Africa, likely in both West and East Africa, before 2500 BCE and by 400 BCE was long established in all the modern major production regions of the Old World, including sub-Saharan Africa, the Mediterranean Basin, India, and Southeast Asia. Further spread occurred as part of the Columbian Exchange, which brought African germplasm to the Caribbean, the southeastern United States, and South America, and Mediterranean germplasm to Cuba, the southwestern United States and Northwest Mexico.

## 1. INTRODUCTION

Cowpea is a diploid (2n = 22), warm season legume which serves as a major source of calories and protein for many people, especially in developing countries. The bulk of cowpea production and consumption is in sub-Saharan Africa, especially in the Sudano-Sahelian Zone (Ousmane Boukar et al., 2019). About 95% of global production reported in FAOSTAT is in West Africa, with Nigeria being the largest producer and consumer of cowpea, producing 3.4 million tonnes in 2017 (FAOSTAT, 2019; Samireddypalle et al., 2017). Other areas of production include Southeast Asia, the Mediterranean Basin, Latin America, and the United States of America. Just over 7.4 million metric tonnes of dry cowpeas were reported worldwide in 2017 (FAOSTAT, 2019), though these numbers do not include Brazil, Ghana, and some other relatively large producers. Most of the production in sub-Saharan Africa is by smallholder farmers in marginal conditions, often as an intercrop with maize, sorghum, or millet (Ehlers & Hall, 1997). Due to its high adaptability to both heat and drought and its association with nitrogen-fixing bacteria, cowpea is a versatile crop (Ousmane Boukar et al., 2019; Ehlers & Hall, 1997).

Almost the entire aerial section of the cowpea plant is edible and regularly consumed. Most commonly, the dry grain is used. The fresh grains are often consumed during the harvest season, and immature pods are eaten as a vegetable, especially in Southeast Asia. In addition, the tender leaves are consumed as a pot herb, mostly in East Africa (Ousmane Boukar et al., 2019). The dry haulms are harvested and sold as fodder for livestock (Samireddypalle et al., 2017). Beyond direct consumption, cowpea provides important agronomic services. As a legume, the plants form root nodules in cooperation with nitrogen-fixing bacteria and are used as green manure (Fatokun et al., 2002; Singh et al., 2017). Spreading varieties are also utilized as cover crops to prevent soil erosion and reduce the incidence of weeds (Wortman & Dawson, 2015).

Cowpea is a valuable species for cultivation as the effects of global climate change become more pronounced. Regions where cowpea is highly cultivated, such as sub-Saharan Africa, overlap with areas predicted to suffer from increased food insecurity due to climate change (Met Office, 2015). Among the expected effects are more extreme weather events, including deeper and longer droughts and increased heat. Cowpea is well-suited for targeted breeding efforts addressing drought and heat tolerance as it produces high yields under terminal drought conditions, is heat resistant, and requires less water than other commonly cultivated legume species (Agbicodo et al., 2009; Ehlers & Hall, 1997).

### Cultural importance of cowpea

Cowpea is an important food in a wide cross-section of cultures. It is commonly a traditional food eaten as part of celebrations of the New Year. In the American South, a dish called “Hoppin’ John” is eaten. This dish generally consists of cowpeas with a black eye pattern, rice, and often bacon. The origin of this tradition is not clearly established, but may have come from Sephardic Jews, who have a similar tradition based on a tractate from the Babylonian Talmud, still part of the modern celebration of the Jewish New Year (September/October; Aviya Amir, personal communication, 2 November 2019). The Aramaic word for cowpea, transliterated as “rubiya,” is bolded in the following quotation.

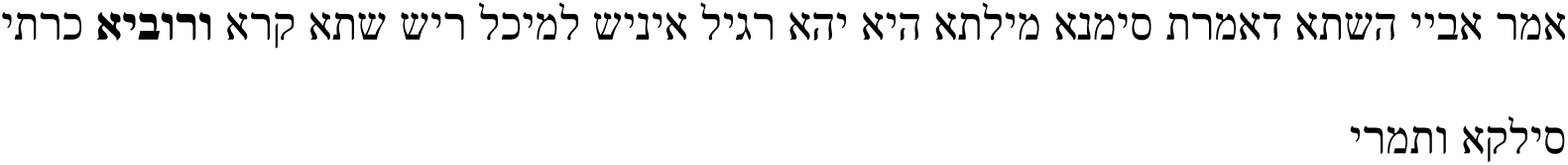

“Abaaye said: Now that you should have a sign, a person should eat at the beginning of the year, a gourd, cowpea, beets, and dates.” (*b.Keritot.6a*)

### Description and nutrition

Cowpea is botanically classified as an annual herb. It demonstrates a wide range of growth habits, ranging from prostrate to erect, can be spreading, climbing, or bushy, and can be determinate or indeterminate. Cowpea is cultivated in a wide range of environments. The specific growth habits of a cultivar or landrace are generally associated with the particular environment and uses. For example, the common black-eye varieties popsular in the United States grow on short bushy plants with determinate growth and with raised racemes on which the pods grow above the foliage, facilitating harvesting. Long bean varieties exhibit climbing and indeterminate growth, allowing for continuous harvesting of green pods over the growing season. Other varieties used as cover crops and green manure to prevent erosion and to fertilize fields are spreading and have indeterminate growth. Some are also day-length sensitive, requiring short days to flower. Some varieties grown in Africa are dual purpose, grown both for seed and animal fodder, providing farmers with additional income, on the order of 25% (Dugje et al, 2009).

Cowpea is grown under a range of cultivation methods. In developed countries, it is mostly grown commercially under irrigation and with fertilizers and applied pesticides, while in developing countries it is mostly grown on smallholder farms as a rainfed subsistence crop, with little to no fertilizer or insecticide input, as well as commonly as an intercrop with maize or other grains, which can lower yield rates. The differences are notable in terms of yield: in the United States the 2017 yield rate for cowpea was 1,700 kg/ha compared to 902 kg/ha in Nigeria and just 464 kg/ha in Uganda (FAOSTAT, 2019).

As a food crop, cowpea is an excellent source of protein, fiber, and a wide range of micronutrients. Cowpea grains are 20-30% protein by dry weight (Ousmane Boukar et al., 2011; Bressani, 1985), and the leaves have a similar protein content (Nielsen et al., 1997). In addition, cowpea is a good source of folic acid, a nutrient of particular importance for pregnant women, and other micronutrients (Ousmane Boukar et al., 2011; Bressani, 1985; Nielsen et al., 1997).

There has historically been some confusion about the taxonomy of cowpea. Historically, cultivated cowpea was separated into a number of different species, including *Vigna catjang, V. sinensis*, and others. Fuller and Murphy (2018) note that cowpea has also been confused with other species, such as horsegram bean, *Macrotyloma uniflorum*. Further confusion has resulted from the transferal of the terms “phaselus” and “phaseolus” in Latin and “frijoles” in some Spanish-speaking countries, which had referred to cowpea, to the New World common bean, *Phaseolus vulgaris* (see Origin and global spread section for more information).

Cowpea taxonomy has been established by Pasquet and Padulosi (2012) as *Dycotyledonea* belonging to the order *Fabales*, family *Fabaceae*, subfamily *Faboideae*, tribe *Phaseoleae*, subtribe *Phaseolinae*, genus *Vigna*, and section *Catjang*. All cultivated cowpea is grouped under *Vigna unguiculata* subspecies *unguiculata*. There are five wild subspecies as well: ssp. *dekindtiana*, ssp. *protracta*, ssp. *pubescens*, ssp. *stenophylla*, and ssp. *tenuis. V. unguiculata* ssp. *dekindtiana* var. *spontanea*, is commonly found throughout sub-Saharan Africa and is believed to be the progenitor of domesticated cowpea (Pasquet and Padulosi 2012).

## 2. MATERIALS AND METHODS

### Genotypic data

Genotypic data was collected from a minicore population consisting of 368 accessions representing worldwide diversity of cultivated cowpea (Muñoz-Amatriaín et al., manuscript in preparation). DNA was extracted from young leaf tissue using the Qiagen DNeasy Plant Mini Kit (Qiagen, Germany). A total of 51,128 SNPs were assayed in each sample using the Illumina Cowpea iSelect Consortium Array (Illumina Inc., California, USA; Muñoz-Amatriaín et al. 2017). Genotyping was performed at the University of Southern California Molecular Genomics Core facility (Los Angeles, California, USA). The same custom cluster file as in Muñoz-Amatriaín et al. (2017) was used for SNP calling..

### Population genetic structure of cultivated cowpea

A total of 42,711 SNPs with minor allele frequencies (MAF) >0.05 in the minicore collection were used for population structure analysis in STRUCTURE 2.3.4 (Pritchard et al., 2000). Detailed information about population genetic structure analyses on this collection can be found in Muñoz-Amatriaín et al. (manuscript in preparation). In brief, STRUCTURE was run for each hypothetical number of subpopulations (*K*) between 1 and 10, and Δ*K* values were calculated according to Evanno et al. (2005) to estimate the optimum number of subpopulations, which was identified as *K*=6. To further understand the relationships between the six subpopulations, information on how those six subpopulations split at different *K* numbers was extracted from STRUCTURE membership assignments for K=1-6 (Figure 1). The geographic origin of most accessions within each subpopulation was established and used for generating Figure 2.

**Figure 1.**
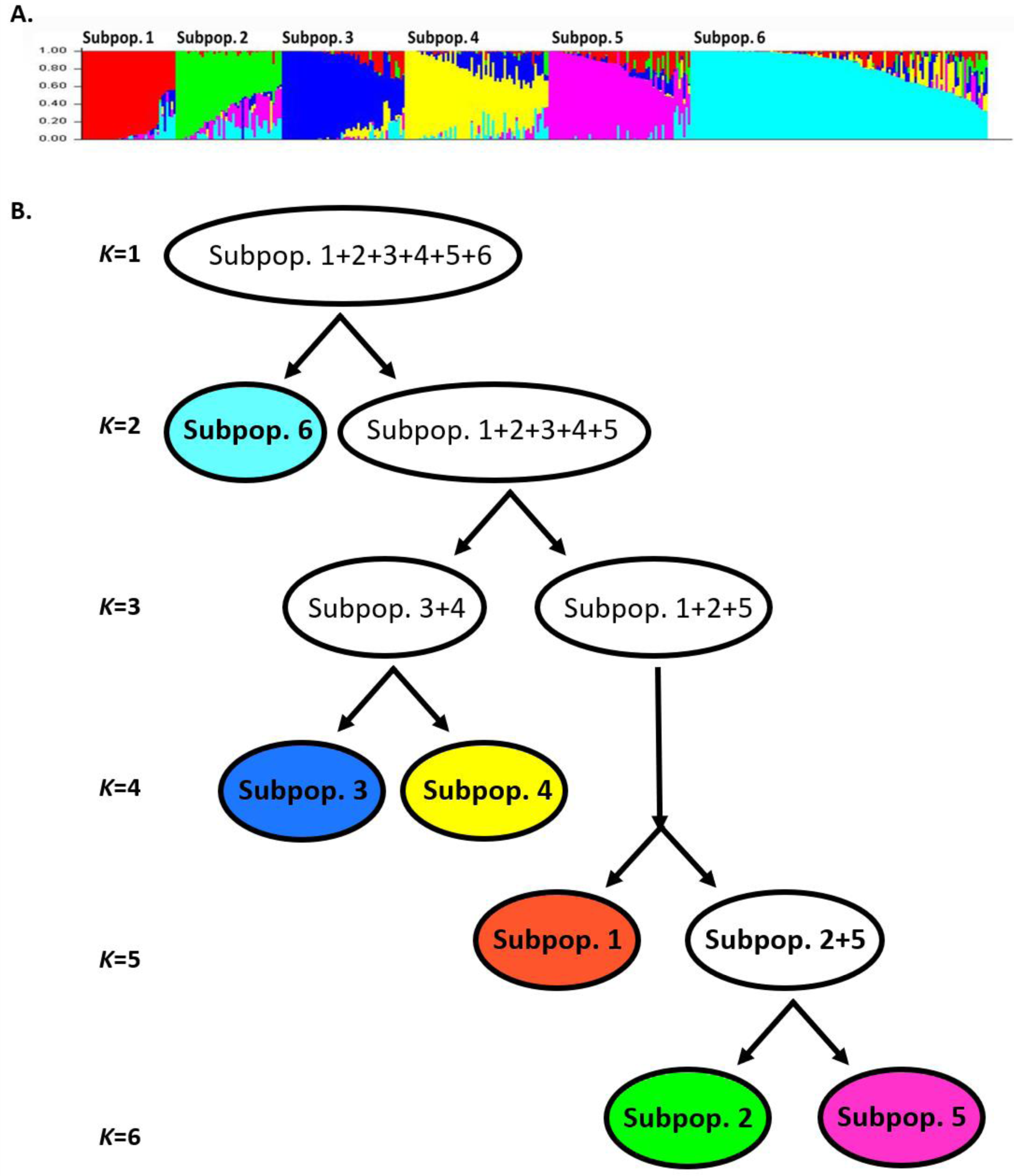
Population genetic structure of cultivated cowpea obtained from STRUCTURE analyses. A) Plot of ancestry estimates for *K*= 6, which each horizontal line representing one accession consisting on *K* colored segments; B) Diagram showing how the six subpopulations divide at different *K* numbers between 1-6.

**Figure 2.**
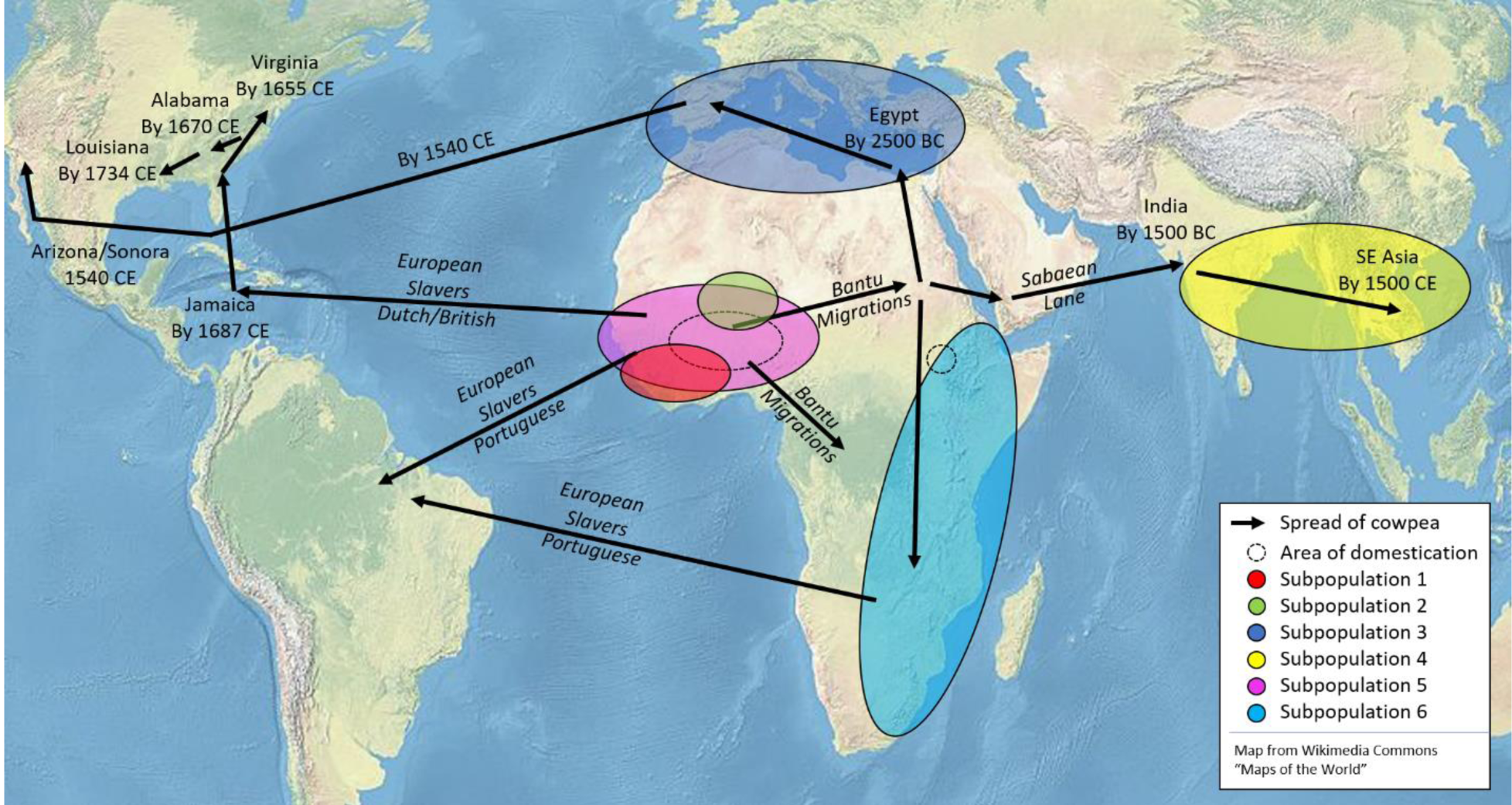
Proposed spread of cowpea from its origins of domestication. Dashed circles indicate likely centers of domestication. Colored circles on the map indicate the geographic area from which most of the minicore accessions assigned to that subpopulation originate. Arrows indicate the proposed routes of spread and are labelled with likely transporters (italicized) or location and date (not italicized).

## 3. Results and Discussion

### Origin and global spread of cowpea

#### Historical linguistics and domestication

Vavilov (1926) was the first to propose a center of domestication for cowpea, proposing China and India as minor centers and Ethiopia as a major center. Later research showed that wild relatives of cowpea are restricted to Africa, ruling out China and India as primary centers of diversity. *V. unguiculata* ssp. *dekindtiana* var. *spontanea* is believed to be the wild progenitor to cultivated cowpea and is found spread over sub-Saharan Africa (Pasquet & Padulosi, 2012). It is a weed, often found on the margins of cultivated fields, which is interfertile with cultivated cowpea, and occurs all over Africa between the Sahara and Kalahari deserts (Coulibaly et al., 2002; Feleke et al., 2006; Rawal, 1975). Most relevant for determining the center of domestication, *spontanea* is known to be particularly interfertile with West African cowpea (Rawal, 1975).

Domesticated cowpea in both West and East Africa shows relatively high levels of diversity compared to other populations worldwide, leading to competing proposals as to whether cowpea was domesticated in West (Ba et al. 2004) or East Africa (Xiong et al., 2018).

Earlier studies using amplified fragment length polymorphism (AFLP; Coulibaly et al., 2002) and ‘random amplification of polymorphic DNA (RAPD) markers (Ba et al., 2004) support domestication in West Africa despite noting that the *Vigna* genus is believed to have evolved in East Africa. The oldest known cowpea in the sub-Saharan African archeological record comes from central Ghana and has been dated to between 1830 and 1595 BCE (D’Andrea et al., 2007). More recently, Huynh et al. (2013) and Xiong et al. (2016) analyzed the population structure of cultivated and wild cowpea using single nucleotide polymorphism (SNP) markers, both showing that cultivated cowpeas in both East and West Africa are most closely related to local wild cowpea. This suggests two independent domestication regions, paralleling the domestication of common bean in both Mesoamerica and the Andean highlands (Kwak et al. 2009), likely with a combining of gene pools, such as is believed to have occurred in Asian rice (Fuller 2011; Vaughan et al. 2008)

Support for the spread of cowpea cultivation out of West Africa comes in the form of linguistic evidence. In the reconstructed proto-Bantu language from circa 3,000 BCE the word for cowpea can be reconstructed as “*-kunde” (Ehret, 1974, 1998; Vansima, 1990). Around this time, Bantu-speaking groups began migrating out from the original Bantu homeland in modern-day Cameroon and Nigeria (Ehret, 1998). From that point forward, it is likely that the word refers to the domesticated plant as wild cowpea is not found in the equatorial rainforest areas that the Bantu moved into. Interestingly, the Bantu-speaking groups which migrated east to the area of modern South Sudan, Sudan, Kenya, and Ethiopia did not bring the term “*-kunde” with them and instead adopted the Southern Cushitic term “*salakw-” (Ehret, 1974, 1998). This could indicate that cowpea cultivation in East Africa predated Bantu arrival and could further support the hypothesis of independent domestication in East Africa, with the two gene pools mixing. The linguistic evidence also rules out domestication in Southern Africa as the common root word for the bean in the area is “*-emba,” which is believed to date to the arrival of Bantu speakers in the region, who likely brought cultivated cowpea with them (Christopher Ehret, personal communication, 7 October 2019). Further, agriculture was not practiced in the region until the arrival of the Bantu speaking groups, making an independent domestication less likely (Blench, 2003).

#### Mediterranean Basin

Cultivated cowpea had spread to Egypt by ∼2500 BCE as evidenced by its presence in royal tombs from the Fifth dynasty, as identified by the prominent German botanist Georg August Schweinfurth (Blench, 2003; El-Din Fahmy, 1997). The Egyptian priesthood had rituals attested to by Plutarch (trans. 1927) in which beans were offered to the gods in the month of Mesore (August-September; *Isis and Osiris*, 378, 68) and that priests avoided eating beans (*Isis and Osiris*, 353, 5). The prohibition on legume consumption is also attested to by Pliny the Elder (trans. 1938) in his Natural History (XVIII.XXX.119). In addition, Pliny notes that the Pythagoreans abstained from eating beans, possibly because they believed the souls of the dead to be contained within (XVIII.XXX.120). It has even been suggested that the prohibition on consuming beans was to avoid farting, which was regarded as impure (Darby et al. 1977).

Cowpea has been known in the Mediterranean Basin since at least the time of the Ancient Greeks. Cowpea was seen as a humble food, both for humans and livestock. In *Georgics*, Virgil writes: “*si uero uiciamque seres uilemque phaselum…*” (“If truly you would sow the lowly kidney bean and vetch…”) (I.227), describing the proper time to sow beans. Traditionally, the word “phaselum” has been translated as “kidney bean” (Virgil trans. 1947). However, “*Phaseolus*” was the genus name assigned to the common bean by Linnaeus, native to the Central and South America, and so Virgil’s “phaselum” can be better understood as simply “bean,” and to specifically refer to cowpea, since during the Roman era, this was the kind of bean commonly grown around the Mediterranean (Albala, 2007). Athenaeus (trans. 1927), in a list of foods, mentions that the Spartans “serve as dessert dried figs, beans, and green calavances” (II.56). The English term calavance (or carauance elsewhere) is an old name for cowpeas and may be a corruption of the Spanish “garbanzo,” referring to *Cicer arietinum*, the chickpea (Wight, 1907). Later, quoting from the lost work *Unhappy Lovers* by Antiphanes: “All the other common desserts are a sign of poverty – boiled chickpeas, beans, apples, and dried figs” (III.101). He even recounts a fart joke regarding a bean-boiling festival in Greece, writing “it is plain that Telemachus constantly fed off pots of beans, and celebrated Bean-Festival as a windy holiday” (IX.407). In each case, Atheneus is using the term “phaselus” as Virgil does.

The authors of antiquity regarded beans as part of a healthy diet. Plato (trans. 1991) writes, in a section discussing what makes a city distinct from a collection of dwellings, that with eating beans “…they will live out their lives in peace and health […] dying as old men…” (II.371d). Galen (trans. 2003) writes about the role of cowpeas in diet twice, calling them both “phaselus” (I.25), as Virgil does, and “dolichos” (I.28) and noting that the whole pod as a vegetable goes by the name “lobio” (I.28), which appears to be the source of the term “lubiya,” the term in modern Arabic, Farsi and Hebrew for cowpea (Wiktionary 2019). Galen mentions cowpeas as part of diet of a man practicing medicine in Alexandria (I.25), likely as a way of indicating that even the simplest of foods could be part of a healthy diet (Albala, 2007).

#### South and Southeast Asia

The earliest definite evidence of cowpea in the Indian sub-continent dates to between 1500 and 1200 BCE at Daimabad, in the Western Zone, with earlier controversial evidence at Hulas, in the Central Zone, dating between 2200 and 1500 BCE (Fuller, 2003). This is roughly contemporary with the earliest confirmed cowpea remains in Africa (D’Andrea et al., 2007). This issue is common with other African domesticates, which often first appear in the archaeological record outside Africa. This may be due to fewer digs being conducted on the continent or simply that fewer archaeobotanical remains have survived, possibly due to specific soil conditions (Blench, 2003; Neumann, 2005). Beans are also generally less likely than cereal grains to survive in the archaeological record due to the way they are prepared and cooked. Beans are generally boiled, and so are less likely to fall into the fire and be carbonized or have the hulls pressed into bricks (Caracuta et al., 2017). Additionally, legumes are known to produce very few phytoliths compared to grains, and so are less likely survive in the archeological record (Caracuta et al., 2017; Tsartsidou et al., 2007)

Blench (2003) proposed three possible routes for how African domesticated cowpea could have arrived in India:

1. From Egypt to the Near East, then across the Iranian plateau towards northwest India.
2. The “Sabaean lane,” through modern Yemen, carried on the yearly monsoonal transports to India.
3. Directly across the open ocean from East Africa to India.

Trade routes across central Asia have long played a role in the movement of goods, technologies, and peoples. Most famous is the Silk Road, which connected China to the Eastern Mediterranean. Population genetic structure data shown in Figure 1 from Muñoz-Amatriaín et al. (manuscript in preparation) show that Mediterranean and Southeast Asian varieties of cowpea are closely related, which could indicate gene flow overland. However, the authors are unaware of any archaeological evidence identifying cowpea in central Asia in a time frame that would support spread via this route and have not attempted to explain the phenomenon in this publication.

The importance of the Sabaean lane in ancient trade is well attested. Sabaea, in modern Yemen, was a major trade hub through which the Mediterranean world accessed products from India and the Far East (Bowen & Albright, 1958). The Romans were highly aware of the region’s importance for the flow of trade and even attempted to capture it at one point. The unknown author of *The Periplus of the Erythraean Sea* relates how a pilot named Hippalus discovered that the locals made use of the monsoonal winds to travel to India (Schoff trans. 1912). Indeed, trade between Rome and India is well-attested, with Roman coinage being found in Indian trading cities on the west coast of the subcontinent (Bowen & Albright, 1958). This is the scenario Blench (2003) identifies as the most likely.

The third proposed route, directly across the Indian Ocean from East Africa is the most unlikely, as regular trade across the Ocean did not begin until the early first millennium CE, long after the arrival of cowpea in India (Sinclair et al. 2012).

The arrival of cowpea in Southeast Asia is much less well-documented. The earliest known reference to cowpea is from the 16^th^ century CE, when it was included in the Ming Dynasty Compendium of Materia Medica (*Bencao Gangmu* 本 草 纲 目), compiled by Li Shizhen (trans. 2003) at the end of the 16^th^ century CE, where it is listed as being effective in treating kidney issues as well as part of a treatment for excessive flatulence. Interestingly, Li Shizhen commented that it was oddly not included in the 3^rd^ century CE Classic of Herbal Medicine (*Shennong Bencao Jing* 神 農 本 草 經). This could indicate that cowpea had not arrived in China by 200 CE or it could simply be that it was not used in traditional medicine at the time.

#### New World

Cowpea was one of the many plants brought to the New World as part of the Columbian Exchange. As noted by Carney (2001), the transportation of foodstuffs was dependent on the movement of people, and so necessarily came alongside systems of social and cultural importance. This is most obvious in the methods of food preparation. African slaves brought to the New World brought along with them their knowledge of how to prepare food, including cooking methods, and these methods have become standard in areas of African settlement. One example of this phenomenon is rice preparation. In West Africa, rice is cooked in water, as opposed to the cooking in fat before boiling as is common in the Mediterranean. This method of food preparation was brought with Africans to the southeastern United States (Carney, 2001). Similarly, in the case of cowpea, the consumption of the whole dry bean boiled together with rice, or cooked separately and combined before serving to create a meal, is common between West Africa and areas settled by Africans during the colonial period, including modern Mexico, the southeastern United States, the Caribbean, and South America.

Cowpea may have been brought to the New World as early as 1500 CE, possibly on the same ships that brought slaves from West Africa (Carrier, 1923). Cowpea was likely included in the food served to slaves on the Middle Passage as part of something called “slabber sauce,” a concoction of vegetables, beans, and often spoiled meat poured over rice (Covey & Eisnach, 2009; Harris, 2011). Other authors make claims that cowpea was brought by the enslaved, secreted away on their persons on the Middle Passage, though those claims have no direct evidence and should not be taken as definitive. These sorts of claims are common in oral traditions of maroon communities, communities of escaped slaves and their descendants, such as the Djuka of French Guiana, who claim that female slaves smuggled rice seeds in their hair (Carney, 2001). If this were the case, this would at most be a minor source of germplasm compared to the more regular imports used to feed slaves on the Middle Passage.

Cowpeas were identified by an English traveler in India, Thomas Herbert, who had traveled there as part of an embassy to Persia in 1634. He wrote about a town: “The people … came aboard us in their small canoes, and sold us for other trifles, Coco-nuts, Mangoes, Iacks, greene Pepper, Carauances or Indian Pease, Hens, Eggs, and Buffols, which because rare are deere” (Herbert, 1634). The reference by Herbert to “Indian Pease [sic]” is significant as he had traveled to the West Indies as well, and so was identifying the beans as the same as those in the West Indies (Carrier, 1923).

Cowpeas were established as a crop in Jamaica by 1687-8 CE, as a book published by Hans Sloane recounts (Sloane, 1707). Sloane visited Jamaica in 1687-8 CE as personal physician to the English governor of the island, Christopher Monck, 2nd Duke of Albemarle. Section XXI discusses cowpeas, called “calavances.” Sloane specifically notes the presence of a black eye and a noticeably sweet taste, a common comment about cowpea in early modern writings. Section I may also refer to cowpea varieties with climbing habits, and Sloane notes that these beans first came to Jamaica from Africa. Sections XII through XX discuss other beans called “Phaseolus,” but it is unclear whether they are different varieties of cowpea or another species altogether (if so, these would likely be common bean, which is native to the New World).

The earliest definitive mention of cowpeas on the North American continent comes from a 1666 CE Virginia law which set the value of various agricultural goods when used as in-kind taxes (Hening, 1823). It seems that the English colonists considered cowpea to be a native crop, indicating its arrival prior to the English colonists. Cowpea is attested to by archaeological findings in the Upper Creek village of Fusihatchee, in modern Alabama, by 1670 CE (Gremillion, 1993). Cowpea was identified in North Carolina around 1700 CE by the Surveyor General of the colony, John Lawson (1714). By 1755 CE cowpea was being grown in Virginia for export to other colonies (Douglass, 1755). Cowpea had spread to French Louisiana by 1734 CE, as attested to by the writings of Antoine-Simon Le Page du Pratz, who lived in Louisiana for sixteen years and among the Natchez for eight of those years. From his writing it is clear that at least some Europeans were aware of the provenance of cowpea in the southeastern modern United States as being ultimately from Africa, which he refers to as “Guinea,” an older term for the west coast of the African continent preserved in some country names, such as Guinea and Guinea-Bissau. Le Page du Pratz writes:

> “The first settlers found in the country French-beans of various colours, particularly red and black, and they have been called beans of forty days, because they require no longer time to grow and to be fit to eat green. The Apalachean [sic] beans are so called because we received them from a nation of the natives of that name. They probably had them from the English of Carolina, whither they had been brought from Guinea. Their stalks spread upon the ground to the length of four or five feet. They are like the other beans, but much smaller, and of a brown colour, having a black ring round the eye, by which they are joined to the shell. These beans boil tender, and have a tolerable relish, but they are sweetish, and somewhat insipid.” (Le Page du Pratz, 1774).

The first known written mention of the term “cowpea” in the English language is from a letter written by Thomas Jefferson to an acquaintance, John Taylor, on October 8, 1797: “I have…received all the good kinds of field pea from England, but I count a great deal more on our southern cowpea. If you wish any of them, I will send you a part,” (Jefferson & Holmes, 2002). The Spanish word “caupí” is believed to be a borrowing of the English term, while the traditional name “frijol” was transferred to the New World common bean (*Phaseolus vulgaris*) in some Spanish-speaking countries (Cubero, 1994).

Cowpea was a base food crop in the American South from the colonial period forward. Black slaves grew cowpea in their vegetable gardens throughout the area (Covey and Eisnach, 2009; Morgan, 1998; Mrozowski et al., 2008; Sutch, 1976). Oral histories of former slaves, collected by the Works Progress Administration during the 1930s, make regular mention of cowpea as part of the diet of the enslaved (Covey & Eisnach, 2009).

Cowpea also has a long history in the southwestern United States. However, when and how cowpea arrived in the area is unclear. The commonly accepted narrative, put forward by Castetter and Bell (1942), is that Eusebio Francisco Kino, an Italian Jesuit missionary in service to the Spanish Crown, brought cowpea with him from Spain along with a variety of other crops in 1683 CE. However, neither Kino (trans. 1919) nor his traveling companion, Juan Mateo Manje (trans. 1954), make mention of any such action in their accounts, merely commenting on the presence of beans in the fields of the missions and the local indigenous peoples. Further, while both authors hailed from cowpea-growing regions in Europe (Kino from Italy and Manje from Spain), when they mention the presence of beans they are generally not specific. In a few cases, Manje specifies the type of bean he is referring to, but it is tepary bean (*Phaseolus acutifolius*), not cowpea. In any case, it appears that the introduction was so long ago that the folk memory of the introduction was forgotten (Castetter & Bell, 1942). All efforts by the authors to obtain any record from the native communities identified in the writings of the Jesuits and Spaniards were unsuccessful. It appears that no records from either the Tohono O’odham or the Pima Yacqui exist regarding cowpea introduction. Some evidence, however, can be found in linguistics. A term for cowpea, common across several related Uto-Aztecan languages, is “yorimuni.” The term can be rendered a number of different ways, including “yori muni,” “yori muuni,” “orimuni,” and others. “Muni” simply means “bean” in general sense (Miller 1996; Robert Valencia Jr., personal communication, 11 September 2019) “Yori” can mean mestizo, Mexican, white, or non-Indigenous people, and generally indicates an “other,” pointing to the introduction of the crop from elsewhere (Miller, 1996; Savor Blog Partners, 2018). Cowpea was certainly established as crop by 1775, when an expedition led by Juan Bautista de Anza travelled up the Colorado River. During that expedition, the travelers were given cowpea by a local chief referred to as Captain Palma (Font, trans. 1930).

The first Spanish explorer to sail up the Colorado River was Hernando de Alcorón in 1540. The available information about Alcorón’s trip is limited to a long letter he sent to the viceroy of New Spain following his return. A full account of the trip was promised, but there is no indication that this report was ever submitted (Elsasser, 1979). In the letter, Alcorón wrote that he “showed them wheat and beans, and other seeds, to see whether they had any of those kinds: but they showed me that they had no knowledge of them, and wondered at all of them.” As other kinds of bean, such as teparary bean, were already in cultivation in the area, Alcorón is likely referring specifically to cowpea (Elsasser, 1979). If so, this would indicate the arrival of cowpea to the southwestern modern United States occurred in 1540 CE, brought by Alcorón. However, Alcorón does not specify whether the beans were brought directly from Spain or had been grown in Spanish New World holdings, though the dominance of common bean in Central America suggests that the cowpea had come from Spain.

#### Dual introduction to the United States

The dual introduction of cowpea to the New World has resulted in great confusion. For example, Perrino et al. (1993) sought to elucidate the spread of cowpea using phenotypic data, expecting American cowpea to match West African varieties due to the slave trade. The study did not distinguish where in the United States various lines came from (i.e. southeastern or southwestern regions). This resulted in confusion as in some cases, the average of the American varieties better matched West African varieties, but in many cases best matched Mediterranean varieties. However, utilizing the Cowpea iSelect Consortium Array (Muñoz-Amatriaín et al. 2017) and the University of California, Riverside minicore (Muñoz-Amatriaín et al., manuscript in preparation), we can show the two distinct introductions of cowpea to the USA using both genetic and textual sources. Further genetic evidence from Carvalho et al. (2017) supports this theory, showing linkages between Cuban and Mediterranean varieties, and between sub-Saharan African and South American varieties.

Additionally, Asian longbean (cultivar group *sesquipedalis*) varieties also came to the United States at some point, but the origin is less clear. They may have been brought by the Spanish from their holdings in East Asia, such as the Philippines, or perhaps Chinese laborers brought it to the southwestern United States while working on the transcontinental railroads. For example, Native Seeds/SEARCH (nativeseeds.org), a seed conservation organization based in Tucson, Arizona, has a variety of longbean which genetically matches Asian varieties, but which was collected from the village of Ahome in Sinaloa, Mexico. Documentation of the origins of the varieties collected in the Native Seeds/SEARCH collection is overall sparse and, in some cases, the seeds obtained from the collection do not have visual characteristics matching the photographs on the seed packets. It is possible that the stories attached to the collection of varieties are inaccurate.

#### Summary

Utilizing the above data gathered from genetic, textual, and archeobotanical sources, a proposal of the spread of cowpea from its origin of domestication can be made. Cowpea had two domestication regions, a major one in West Africa and one in Eastern Africa. From the West African domestication, cowpea was spread by the Bantu migrations south into the equatorial rainforest and east across the Sahelian zone to the area of modern Sudan, South Sudan, and Ethiopia. From there, three branches emerged. One branch led down to southern Africa, which merged with the East African domestication. Another branch moved north, likely up the Nile, to Egypt, where it was present by 2500 BCE, and then spread across the Mediterranean Basin, where it was established enough to be considered a basic food crop by 400 BCE. This Mediterranean population was brought to Spain’s colonial holdings in the New World, including to the modern southwestern United States. The third branch went east, most probably via the “Sabaean lane” in modern Yemen, from which it reached the west coast of India by 1500 BCE, and from there spread to Southeast Asia, where the *sesquipedalis* cultivars were selected for. During the 16^th^ century CE, cowpea was brought from West Africa to the New World, mostly on slaving ships to colonial slave societies, including modern Brazil, the Caribbean, and the American South. At the same time, Spanish colonists and explorers brought Iberian cowpea to the southwestern United States and northwestern Mexico. An analysis of population genetic structure of cultivated cowpea can be found in Figure 1 and a map showing the proposed spread can be found in Figure 2.

Similar proposals regarding the spread of cowpea have been made previously, most notably by Steele and Mehra (1980) and Ng and Maréchal (1985). However, such older proposals lack the depth of evidence to support their claims and instead make conjectures with more limited data. For example, both above publications contend that there was a spread of cowpea from India towards the Mediterranean through the Iranian plateau, but this is not borne out by available archaeological data. Additionally, these publications put the date of arrival to India around 150 BCE and to the Mediterranean Basin around 300 BCE, far later than the archaeological evidence attests.

## DATA AVAILABILITY

The genotypic data on the minicore that support the findings of this study will be provided in a publication by María Muñoz-Amatriaín et al.

## ACKNOWLEDGEMENTS

The information contained in this manuscript is also published in the Ph.D. thesis of author I.A. Herniter, “Genetics of Consumer-Related Traits in Cowpea (*Vigna unguiculata* [L.] Walp.); the authors thank: Sassoum Lo, Yi-Ning Guo, and Yoko Hiraoka for helpful discussion; Aviya Amir for assistance regarding the cultural importance of cowpea in the Jewish tradition; Christopher Ehret for explaining the African linguistic evidence. This study was supported by the Feed the Future Innovation Lab for Climate Resilient Cowpea (USAID Cooperative Agreement AID-OAA-A-13-00070), the National Science Foundation BREAD project “Advancing the Cowpea Genome for Food Security” (NSF IOS-1543963) and Hatch Project CA-R-BPS-5306-H.

